# A versatile reporter system to monitor virus infected cells and its application to dengue virus and SARS-CoV-2

**DOI:** 10.1101/2020.08.31.276683

**Authors:** Felix Pahmeier, Christoper J Neufeldt, Berati Cerikan, Vibhu Prasad, Costantin Pape, Vibor Laketa, Alessia Ruggieri, Ralf Bartenschlager, Mirko Cortese

## Abstract

Positive-strand RNA viruses have been the etiological agents in several major disease outbreaks over the last few decades. Examples of that are flaviviruses, such as dengue virus and Zika virus that cause millions of yearly infections and spread around the globe, and coronaviruses, such as SARS-CoV-2, which is the cause of the current pandemic. The severity of outbreaks caused by these viruses stresses the importance of virology research in determining mechanisms to limit virus spread and to curb disease severity. Such studies require molecular tools to decipher virus-host interactions and to develop effective interventions. Here, we describe the generation and characterization of a reporter system to visualize dengue virus and SARS-CoV-2 replication in live cells. The system is based on viral protease activity causing cleavage and nuclear translocation of an engineered fluorescent protein that is expressed in the infected cells. We show the suitability of the system for live cell imaging and visualization of single infected cells as well as for screening and testing of antiviral compounds. Given the modular building blocks, the system is easy to manipulate and can be adapted to any virus encoding a protease, thus offering a high degree of flexibility.

**IMPORTANCE:** Reporter systems are useful tools for fast and quantitative visualization of viral replication and spread within a host cell population. Here we describe a reporter system that takes advantage of virus-encoded proteases that are expressed in infected cells to cleave an ER-anchored fluorescent protein fused to a nuclear localization sequence. Upon cleavage, the fluorescent protein translocates to the nucleus, allowing for rapid detection of the infected cells. Using this system, we demonstrate reliable reporting activity for two major human pathogens from the *Flaviviridae* and the *Coronaviridae* families: dengue virus and SARS-CoV-2. We apply this reporter system to live cell imaging and use it for proof-of-concept to validate antiviral activity of a nucleoside analogue. This reporter system is not only an invaluable tool for the characterization of viral replication, but also for the discovery and development of antivirals that are urgently needed to halt the spread of these viruses.

## INTRODUCTION

Positive sense single stranded RNA viruses constitute a major fraction of endemic and emerging human viruses (1). Among the positive-strand RNA viruses, flaviviruses such as dengue virus (DENV) and Zika virus (ZIKV) are some of the most prevalent arboviral pathogens and are considered a major public health problem (2, 3). Currently, there are no universal vaccines or specific antiviral drug approved for the prevention or treatment of infections with these viruses (4). Members of the *Coronaviridae* family also have a positive-strand RNA genome and have caused major outbreaks in the last two decades (5, 6). Currently, the world is facing the pandemic outbreak of SARS-CoV-2, the causative agent of coronavirus disease 2019 (COVID-19) (7, 8). As of August 2020, over 19 million confirmed cases and more than 700,000 confirmed deaths have been reported in 216 countries (9). Despite immense efforts by research teams around the world, there is still a dire need for effective and widely available treatment options and a prophylactic vaccine.

Once released into the cell, the full genome of flaviviruses and the large open reading frame (ORF1ab) of coronaviruses are translated as polyproteins. Signal peptides and internal transmembrane regions direct polyprotein synthesis to the endoplasmic reticulum (ER) membrane where co-translational cleavage generates the mature viral proteins (10, 11). The flaviviral protease NS2B/3, together with host proteases, cleaves the flavivirus polyprotein into three structural and seven non-structural proteins (12, 13). In the case of coronaviruses, ORF1ab is expressed as two polyproteins, which are cleaved into sixteen non-structural proteins (nsp) by the viral papain-like protease (PL_pro_) residing in nsp3 and the 3C-like protease (3CL_pro_) of nsp5 (14–17). The replication of viral RNA of both virus groups was shown to occur on ER derived membranes, in specialized virus-induced membrane compartments termed replication organelles (10, 11, 18–20).

Reporter systems for detection of virus infection are an invaluable tool for the characterization and quantification of virus infection kinetics, for the characterization of virus-host cell interactions and for the identification of antiviral compounds. One approach is the insertion of tags into the viral genome that, upon replication and translation, allow for visualization of the infected cells. However, this approach requires functional molecular clones of a given genome, which are not always available. In addition, insertion of a tag frequently causes attenuation of viral replication competency and therefore, the search for adequate insertion sites is time-consuming or might fail.

An alternative approach is the use of engineered fluorescent reporter proteins stably expressed in cells and altering their subcellular distribution upon viral infection (21–23). Building on this idea, here we established a reporter system based on an ER-anchored green fluorescent protein (GFP) that upon cleavage by a viral protease is released from the ER and translocated into the nucleus. Using this system, we demonstrate the reliable reporting activity of DENV and SARS-CoV-2 infected cells. Moreover, we apply this reporter cell system to live cell imaging and assessment of an antiviral compound.

## MATERIALS AND METHODS

### Cell lines and virus strains

HEK-293T, A549 and VeroE6 cells were purchased from ATCC; Huh7 cells (24) were obtained from Heinz Schaller (Center for Molecular Biology, Heidelberg). Generation of the cell lines Huh7-Lunet and the derivative Huh7-Lunet-T7, stably expressing the RNA polymerase of bacteriophage T7, have been previously described (25, 26). All cells were cultured at 37°C and 5% CO_2_ in Dulbecco’s modified Eagle medium (DMEM, Life Technologies) containing 10% fetal bovine serum, 100 U/mL penicillin, 100 μg/mL streptomycin and 1% non-essential amino acids (complete medium). Huh7-Lunet-T7 cells were cultured in complete medium, supplemented with 5 μg/mL zeocin. A549-ACE2 were generated by transduction of A549 with lentiviruses encoding for the human Angiotensin-converting enzyme 2 (ACE2) gene as previously described (41).

Wild-type (WT) DENV-2 was produced from an infectious molecular clone based on strain 16681 as described elsewhere (27). The DENV reporter virus genome encoding the Turbo far red fluorescent protein FP635 (DENV-faR) has been previously described (28). SARS-CoV-2 (strain BavPat1) was kindly provided by Prof. Christian Drosten (Charité Berlin, Germany) through the European Virology Archive. Except for DENV-faR that was generated by electroporation of BHK-21 cells as previously described (28), all virus stocks were generated by infection of VeroE6 cells. Supernatants were harvested, filtered, and virus concentration was determined by plaque assay. For infection experiments, cells were inoculated as specified in the results section for 1 h at 37°C. Fresh complete medium was then added, and cells were incubated for the indicated time spans.

### Antibodies

The antibodies used in this study are listed in Table 1.

**TABLE 1.** List of antibodies used in this study.

| Antibodies | Concentration |  | Source |
| --- | --- | --- | --- |
|  | WB | IF |  |
| Mouse IgG1 anti-DENV NS3<sup>a</sup> | 1:1000 | 1:200 | GeneTex |
| Mouse anti-GAPDH<sup>a</sup> | 1:1000 | n.a. | Santa Cruz Biotechnology |
| Rabbit anti-GFP<sup>a</sup> | 1:1000 | n.a. | Roche |
| Mouse IgG2a anti-dsRNA<sup>a</sup> | n.a. | 1:400 | Scicons |
| Mouse IgG1 anti-SARS-CoV N<sup>a</sup> | n.a. | 1:500 | Sino biologicals |
| Goat anti-mouse IgG-HRP<sup>b</sup> | 1:10000 | n.a. | Sigma-Aldrich |
| Goat anti-rabbit IgG-HRP<sup>b</sup> | 1:10000 | n.a. | Sigma-Aldrich |
| Alexa Fluor 568 donkey anti-mouse IgG<sup>b</sup> | n.a. | 1:1000 | Thermo Fisher Scientific |
| Alexa Fluor 568 donkey anti-rabbit IgG<sup>b</sup> | n.a. | 1:1000 | Thermo Fisher Scientific |

a: primary antibody, b: secondary antibody; WB: western blot; IF: immunofluorescence.

### Generation of the reporter construct

A synthetic DNA construct containing the sequence encoding the reporter protein was generated by Integrated DNA technologies (Coralville, IA, USA). The reporter sequence was inserted into the lentiviral vector pWPI via AscI and SpeI restriction sites (pWPI-RC). Oligonucleotides encoding the protease cleavage sites were designed to allow insertion into the vector via MluI and BamHI restriction sites. The primer pairs (Table 2) spanning a given protease cleavage site were heated to 95°C and allowed to anneal by decreasing the temperature in 5°C increments every 2 min. The resulting double-stranded DNA product was inserted via MluI and BamHI into pWPI-RC and ligation products were amplified in *E. coli* (strain DH5α). Integrity of amplified plasmids was confirmed by restriction pattern analysis and sequence analysis of the insert region, respectively. The complete nucleotide and amino acid sequences of the reporter construct are available on request. The expression of the reporter construct was under the control of the eukaryotic translation elongation factor 1-alpha promoter.

**TABLE 2.**
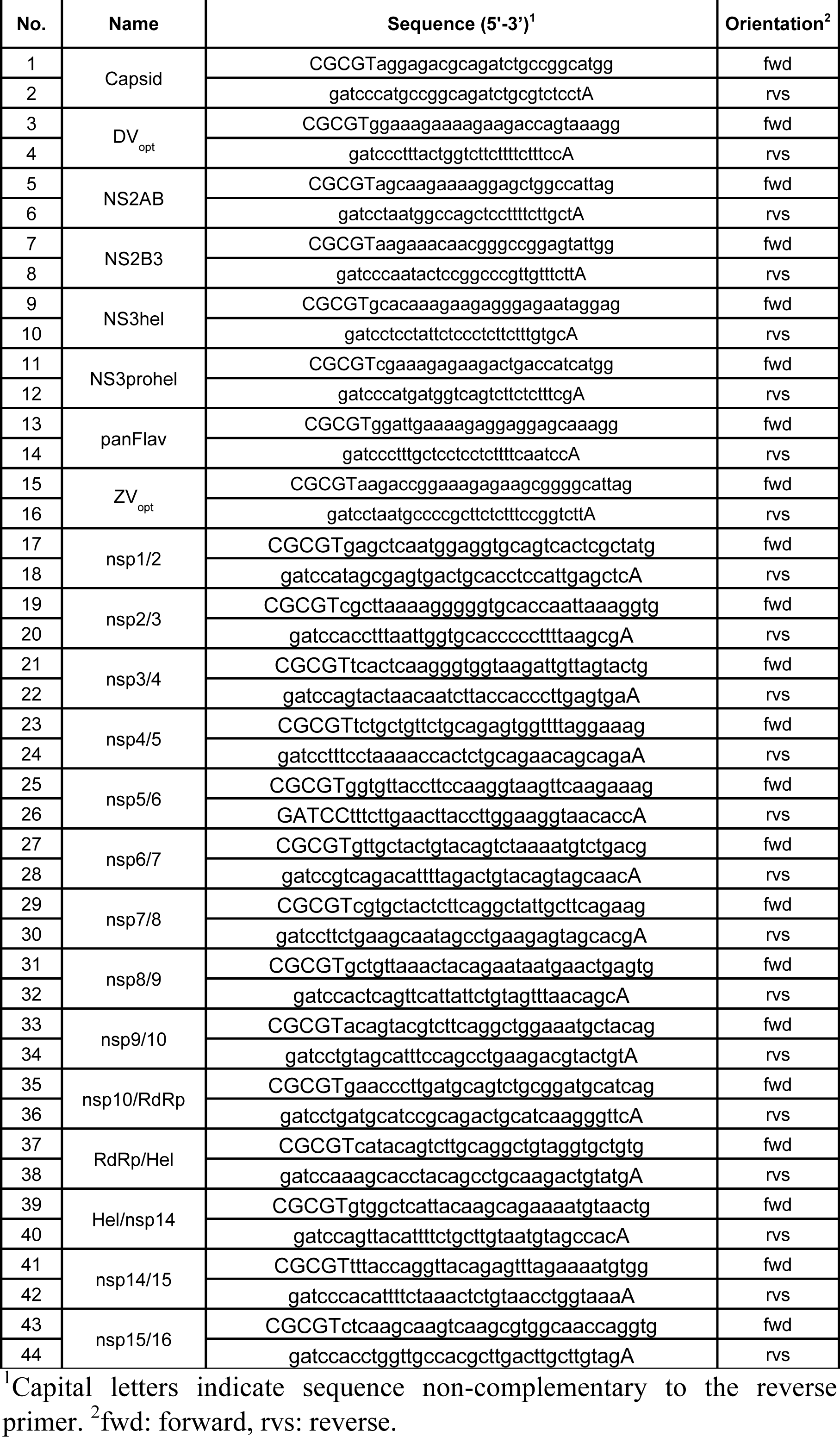
Sequences of oligonucleotides used in this study.

### Lentiviral transduction and generation of reporter cell lines

Cells stably expressing the protease reporter constructs were generated by lentiviral transduction. Subconfluent HEK-293T cells were transfected with the pWPI vector encoding the reporter construct together with packaging plasmids pCMV-Gag-Pol and pMD2-VSV-G (kind gifts from D. Trono, EPFL, Lausanne). After 2 days, the supernatant of transfected cells was harvested, filtered, and stored at −80°C. Lentiviruses were titrated by SYBR green I-based real-time PCR-enhanced reverse transcriptase (SG-PERT) assay (29, 30) using the Takyon SYBR green kit (Eurogentec). The titer was determined by comparison with a standard curve of known RNA concentrations. Lentiviral transduction was performed by addition of the filtered supernatant to Huh7, Huh7-Lunet-T7 or A549-ACE2 cells (multiplicity of infection (MOI) = 5) in presence of 4 μg/mL of polybrene. For the generation of stable cell lines expressing the reporter constructs, cells were cultured in medium containing 1 μg/mL puromycin. Cells stably expressing the SARS-CoV-2 optimized reporter construct were FACS-sorted to obtain single cell clones with homogenous expression levels of the fluorescent reporter.

### Indirect immunofluorescence (IF)

Cells were seeded on glass cover slips and harvested at the indicated time points. The cells were washed once with PBS and fixed with paraformaldehyde (PFA, 4% in PBS) at room temperature (RT). PFA was removed, cells were washed once with PBS and the cover slips were covered with PBS containing Triton X-100 (0.2%) to permeabilize the cells. Cells on cover slips were blocked with skimmed milk (2%) in PBS containing Tween20 (PBS-T (0.02%)) for 1 h. After blocking, the cover slips were placed on 30 μL of primary antibody, diluted in the blocking buffer, with the cell side facing the drop. Cells were incubated for 1.5 h at RT and washed thrice with PBS-T (0.02%). The cover slips were then placed on 30 μL of secondary antibody with the cell side facing the drop. After 45 min of incubation at RT, the cells were washed thrice with PBS-T (0.02%) and cover slips were mounted on microscopy slides using Dapi-Fluoromount-G mounting media (Southern BioTech).

### Western blot

Cells were washed once with PBS and lysed in western blot lysis buffer (1% Triton X-100). After sonification and denaturation at 95°C, protein concentration was measured by Bradford assay. Cell lysates were mixed with Bradford reagent (1:5; Bio-Rad) and absorbance was measured at 595 nm. For each sample, 10 μg of total lysate was resolved by electrophoresis into a 10% or 15% sodium dodecyl sulfate-polyacrylamide gel for NS3 or GFP, respectively. Proteins were transferred to a polyvinylidene fluoride membrane overnight at 4°C. Membranes were blocked in skimmed milk (5%) in PBS-T (0.2%) for 1 h at RT. After washing thrice with PBS-T (0.2%) for 15 min, membranes were incubated with primary antibody for 1 h at RT. The membranes were washed thrice and HRP-conjugated secondary antibody was added. After incubation for 1 h at RT, the membrane was washed thrice, and the bound antibodies were detected using enhanced chemiluminescence solution (Perkin Elmer, Waltham, MA, USA). Images were acquired using the ChemoCam 6.0 ECL system (INTAS Science Imaging, Goettingen, Germany).

### Live cell imaging

Huh7-Lunet-T7 cells expressing the dengue reporter constructs (Lunet-T7-RC) were seeded onto a glass bottom 35 cm^2^ dish (Mattek) at a density of 2 × 10^4^. Transfection of the pIRO-D system (40) was performed 24 h post-seeding using TransIT-LT1 (Mirus Bio) transfection reagent according to the manufacturer’s instructions. Four hours post transfection (hpt) the transfection medium was exchanged for complete medium lacking phenol red (imaging medium). Images were collected with a Perkin Elmer spinning disk confocal microscope. For SARS-CoV-2 live cell imaging, A549-ACE2 or a selected clone of A549-ACE2 stably expressing the fluorescent reporter (A549-ACE2-RC), were seeded on 35 mm dishes (Ibidi) with gas permeable membrane and sealable lid. Cells were infected for 1 h with SARS-CoV-2 (MOI = 10) and 2 hpi the medium was exchanged for imaging medium. Lid was moved to the locked position and silicon was used to seal the dish in order to prevent evaporation. Images were collected with a 20x lambda air objective on a Nikon Eclipse Ti widefield microscope. Multiple observation fields were imaged for 8 h or 18 h at an interval of 10 min for transfection or infection, respectively.

### Compound screening assay

A549-ACE2 cell clones were seeded in duplicates for each condition. On the next day, the cells were treated with a serial dilution of 1:3, starting at 1.1 μM Remdesivir (Hoelzel-biotech, Germany) or with the solvent dimethyl sulfoxide (DMSO) serving as a control. After 30 min, cells were infected with SARS-CoV-2 (MOI = 5) in the presence of the compound and 16 h later, cells were fixed and stained for N protein by IF. Images were acquired with a Perkin Elmer spinning disk confocal microscope. Signal intensity of N protein staining and nuclear GFP signal was quantified on a single cell level by a semi-automated image analysis workflow (31). Cells were considered as positive for infection in the reporter cell line when the nuclear GFP signal intensity was greater than 7,000 arbitrary fluorescence units. Inhibition was quantified by normalizing the values to those obtained with cells that were treated with DMSO only (no inhibition).

### Bioinformatics analysis

Images were analyzed using the Fiji software (32, 33). Graph generation and statistical analysis was performed using the GraphPad Prism 8.1 software package. The scheme of the assumed reporter topology was designed with the Illustrate software (34) using the RCSD PDB entries 4EVL, 4RXH (chain A), 4CG5 (chain C) and 2VBC (chain B).

## RESULTS

### Design and characterization of DENV reporter constructs

In order to generate a reporter system that can specifically indicate virus infection, we designed a construct expressing a GFP fusion protein that could selectively be cleaved by viral proteases. The reporter construct was engineered for viruses that produce ER tethered polyproteins that are processed by viral proteases in close proximity to ER membranes. The transmembrane (TM) domain of the ER-resident protein sec61β was used to target the reporter protein to ER membranes (Figure 1). This ER anchor was connected to a GFP moiety containing the nuclear localization signal (NLS) sequence from simian virus 40 large T-Antigen via a variable linker. The linker region was flanked by restriction enzyme recognition sites allowing the easy insertion and screening of different protease cleavage sequences (Figure 1A). Protease cleavage of the linker would result in GFP translocation from the cytosolic ER to the nucleus, which can be easily detected and quantified by light microscopy.

**Figure 1:**
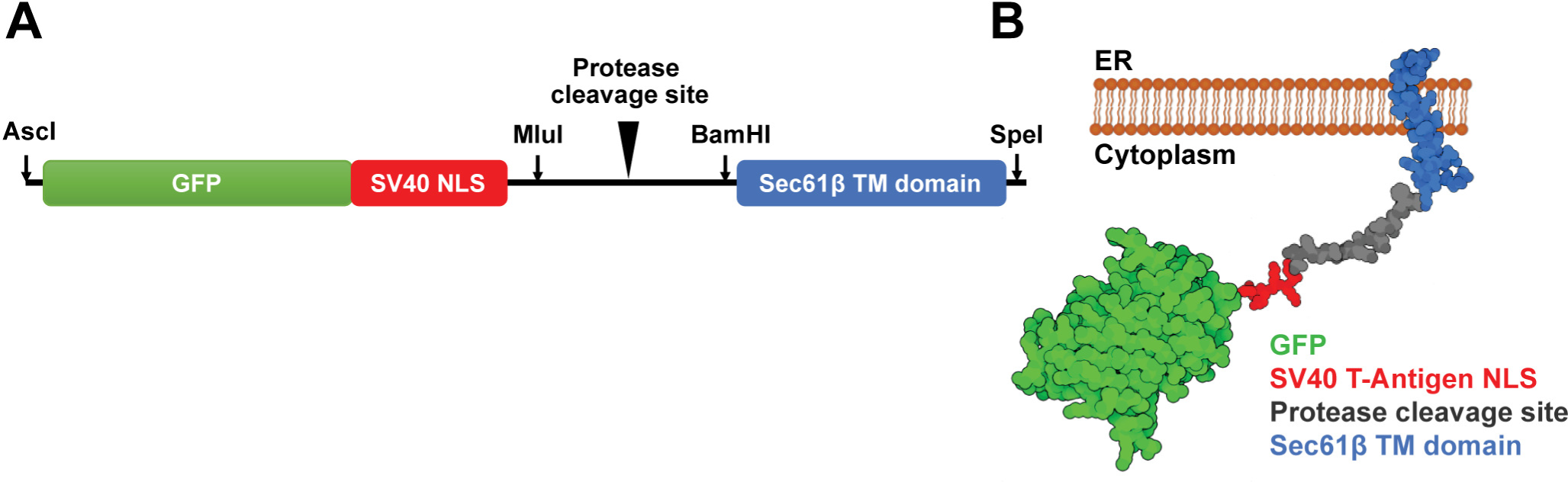
Schematics of the reporter construct (A) and the predicted membrane topology (B). A) Arrows indicate restriction sites for MluI and BamHI that flank the linker region and allow the insertion of the protease cleavage site. B) Proteins and peptides are colored as indicated on the bottom right of the panel.

The DENV polyprotein is cleaved into the individual viral proteins by either the host signal peptidase of the viral NS2B/3 serine protease (12, 13). The ER-resident NS2B protein acts as a co-factor of NS3 protease and anchors it to ER membranes (35, 36). To determine an optimal system for reporting DENV infection, several previously described NS2B/3 specific cleavage sequences were inserted into the reporter construct (Table 3).

**TABLE 3.** List of DENV cleavage site sequences inserted into the reporter construct.

| No. | Name | Cleavage site sequence<sup>1</sup> | Source |
| --- | --- | --- | --- |
| 1 | Capsid | RRRR↓SAGM | Shiryaev *et. al.* 2007 (37) |
| 2 | DV<sub>opt</sub> | GKKRR↓PVK | Shiryaev *et. al.* 2007 (37) |
| 3 | NS2AB | SKKR↓SWPL | Shiryaev *et. al.* 2007 (37) |
| 4 | NS2B3 | KKQR↓AGVL | Shiryaev *et. al.* 2007 (37) |
| 5 | NS3hel | AQRR↓RRIG | Shiryaev *et. al.* 2007 (37) |
| 6 | NS3prohel | RKRR↓LTIM | Shiryaev *et. al.* 2007 (37) |
| 7 | panFlav | GLKR↓GGAK | Shiryaev *et. al.* 2007 (37) |
| 8 | ZV<sub>opt</sub> | KTGKR↓SGAL | Arias-Arias *et. al.* 2020 (38) |

<sup>1</sup> Cleavage site is indicated with ↓.

Reporting activity of the cells expressing each of the constructs was first tested by assessing the subcellular GFP localization by IF. Reporter cell lines were generated by lentiviral transduction of Huh7 cells at an MOI of 5 to ensure maximal transduction efficiency. Subsequently, the cell pools expressing an individual construct were infected with DENV for 48 h (MOI = 5) to observe GFP localization in infected versus non-infected cells. To ensure specificity of GFP translocation, cells were fixed and stained for DENV NS3 protein.

In DENV-infected cells, NS3 was observed in the perinuclear region as previously described (10, 18, 39). While the GFP signal from the reporter constructs 1, 2, 3 and 7 showed an ER-like pattern in mock infected cells, nuclear signal was observed in cells expressing constructs 4, 5, 6 and 8 already in the absence of DENV (Figures 2A and 2B). In infected cells expressing the reporter construct 1, a noticeable increase of nuclear GFP signal was found (Figure 2C). Increased nuclear GFP localization, upon DENV infection, was also observed in reporter cell lines expressing constructs 3, 4, 5, 7, and 8. No clear differences of GFP localizations were seen for constructs 2 and 6 following virus infection. For construct 1, all cells with nuclear GFP signal were also positive for NS3 staining and an additional ∼6% of cells were positive for NS3 staining alone (Figures 2C and 2D). The other constructs showed a higher percentage of cells only positive for NS3 staining and in cells expressing constructs 2, 6 and 7, nuclear GFP signal was observed in the absence of NS3 staining (Figure 2C).

**Figure 2:**
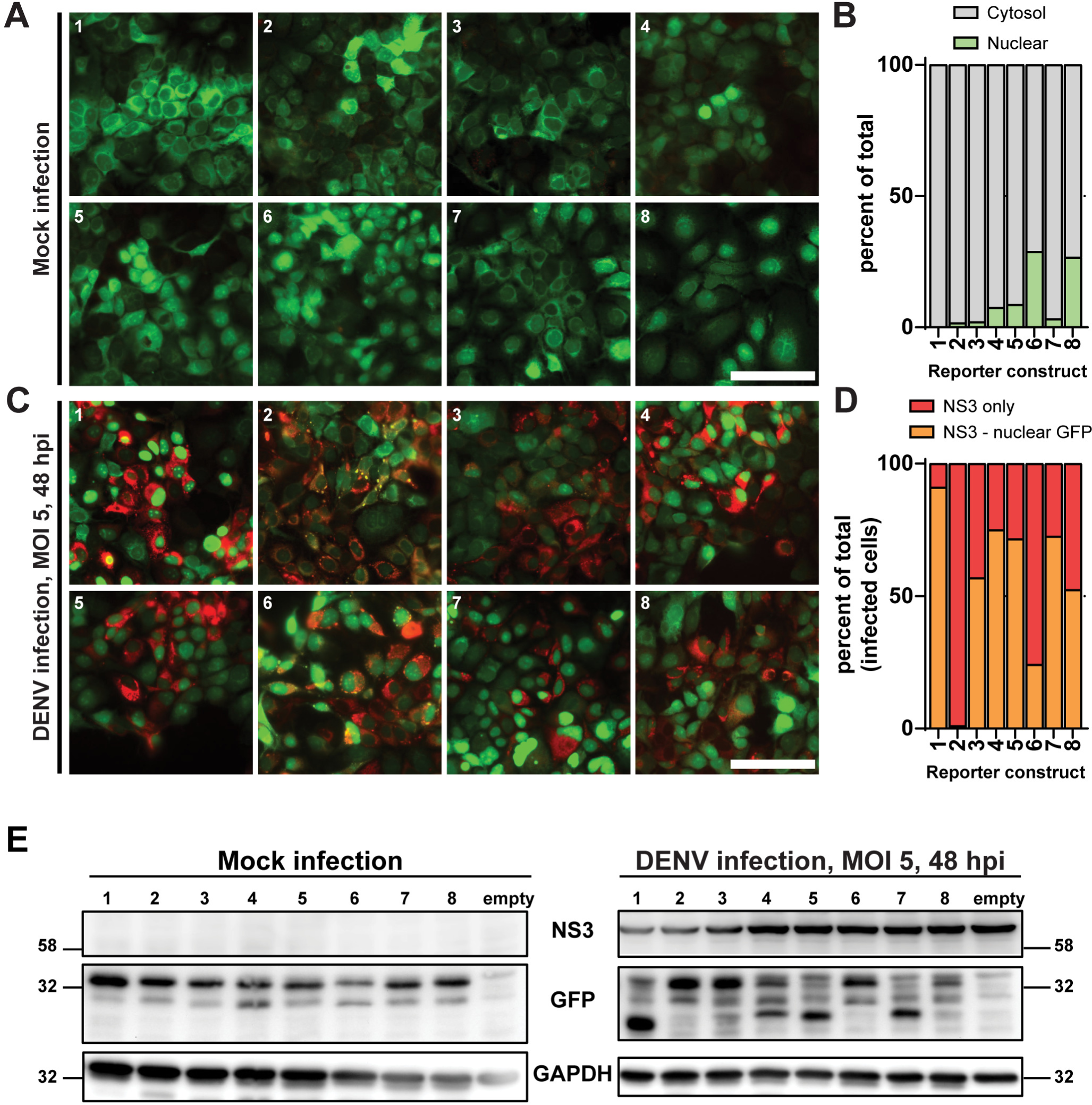
Evaluation of DENV reporter constructs. A) Huh7 cells were transduced with lentiviruses encoding for the different DENV GFP-based reporter constructs 1-8 (Table 3) at an MOI of 5. Cells were fixed 72 hours post-transduction and subcellular localization of GFP was analyzed by confocal microscopy. Scale bar: 100 μm. B) Quantification of images as in A). The percent of cells showing nuclear or cytosolic GFP localization is shown. C) Huh7 cells were transduced as above for 24 h before being infected with DENV at an MOI of 5. Cells were fixed 48 hpi and NS3 was stained by immunofluorescence. Subcellular localization of NS3 and GFP were analyzed by confocal microscopy. Red: DENV NS3 protein; green: reporter GFP signal. Scale bar: 100 μm. D) Quantification of images as in C). Percent of cells positive for NS3 (red) and cells positive for both nuclear GFP and NS3 signals was quantified. E) Cells expressing the reporter constructs 1-8 or an empty plasmid (empty) were infected with DENV (MOI = 5). 48 hpi cells were lysed and 10 μg of total protein for each sample was resolved by SDS-PAGE. NS3 and GFP were detected with a specific antibody. Glyceraldehyde-3-phosphate dehydrogenase (GADPH) served as a loading control. Size of the pre-stained protein ladder bands is indicated in kDa on the side of each panel.

To investigate cleavage of the various reporter proteins in DENV infected cells, we used western blot analysis of cell lysates prepared 48 h post infection (Figure 2E). Based on the construct design, the cleaved GFP-NLS fusion protein was predicted to have a molecular weight of ∼34 kDa. In the mock infected cell lines, we observed a GFP protein with the expected molecular weight and additional bands that were also found in cells transfected with empty plasmid. Upon DENV infection, additional ∼30 kDa GFP-positive bands were detectable in cell lysates containing reporter constructs 1, 4, 5, 7 and to a lesser extent in cell lysates with constructs 3 and 8, which is consistent with the predicted cleavage product size. Notably, construct 1 showed higher levels of specific cleaved product compared to the other cell lines.

To summarize, DENV infected cells expressing reporter construct 1, containing the NS2B/3 cleavage site from the capsid region, showed the highest level of the correct cleavage product in western blot analysis as well as robust and specific nuclear translocation of GFP signal by fluorescence microscopy. Therefore, reporter construct 1 was chosen for generation of stable cell lines for further characterization. These Huh7-derived cell lines were designated reporter construct (RC) cell lines Huh7-RC and Lunet-T7-RC.

### Time-course experiments confirm early and reliable identification of DENV NS2B/3 positive cells

Next, we analyzed the kinetics of GFP translocation in the reporter cell line Huh7-RC. Cells were infected with either the WT DENV or with a DENV reporter expressing a far red fluorescent protein (DENV-faR; MOI = 5) (28). Cells were fixed at 24 h, 48 h or 72 h post infection and subsequently analyzed by wide-field fluorescence microscopy (Figure 3A).

**Figure 3:**
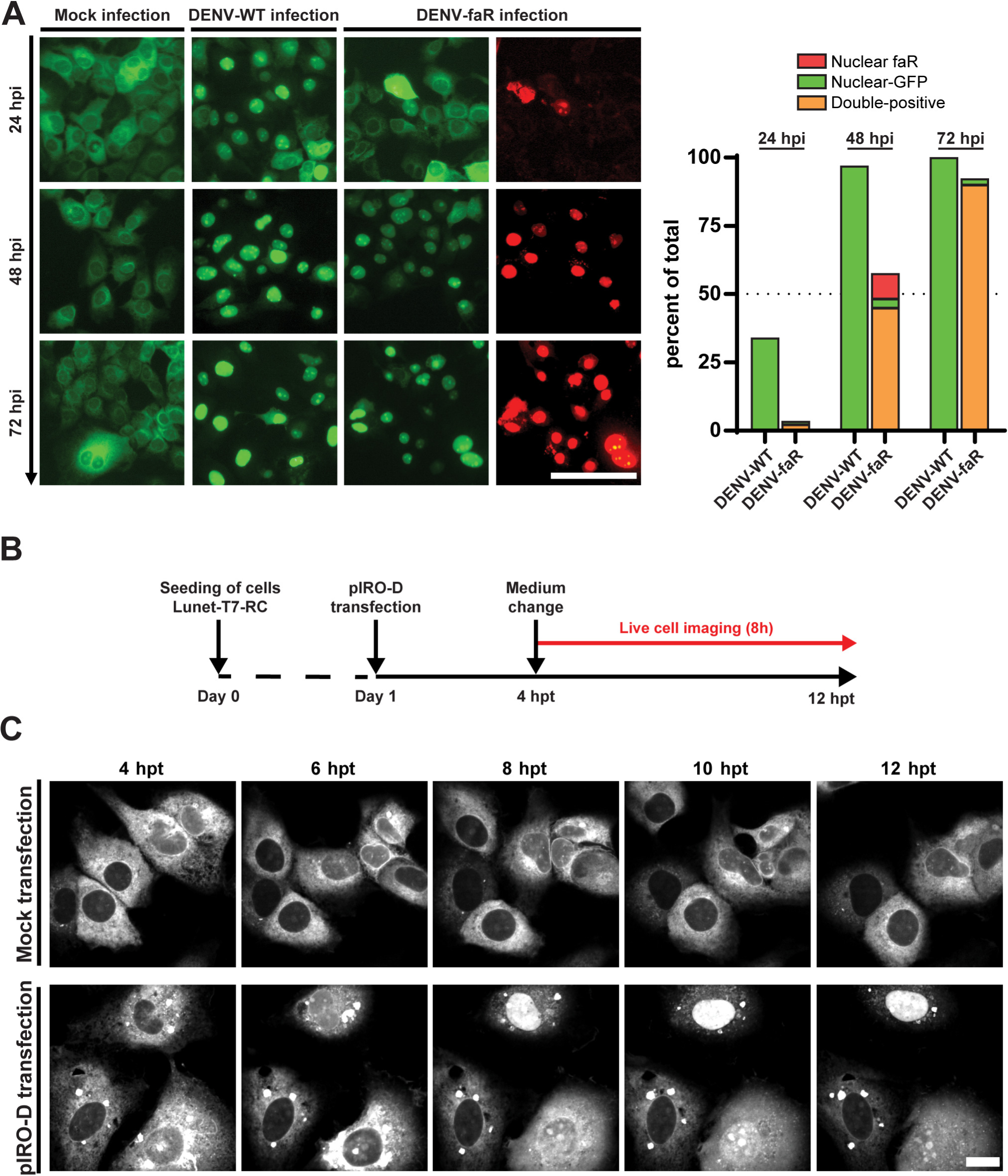
Time-course experiments of DENV reporter in infection and transfection systems. A) Huh7 cells stable expressing the reporter construct 1 were mock infected, infected with DENV WT or the reporter virus DENV-faR (MOI = 5). Left panel: Cells were fixed at the indicated hpi and signals of the reporter virus (red) and the GFP-based reporter construct (green) were detected with a wide-field fluorescence microscope. Scale bar: 100 μm. Right panel: quantification of images in the left panel. Percentage of cells positive for nuclear GFP signal (Nuclear-GFP), DENV-faR reporter virus (Nuclear-faR) and positive for both nuclear GFP and faR reporter signals (orange) was quantified. Values were normalized by setting the total number of cells counted using DAPI staining as 100%. B) Experimental set-up to monitor GFP-reporter activation in pIRO-D transfected cells. C) Lunet-T7-RC cells stably expressing the T7 RNA polymerase and the reporter construct 1 were mock or pIRO-D transfected. 4 hpt the medium was changed, and live cell imaging performed for 10 h with a confocal spinning disc microscope. Images of representative fields of view and the indicated time points are shown. Scale bar: 20 μm.

Mock infected reporter cells exhibited the predicted ER-like localization of the GFP signal. In contrast, reporter cells infected with WT virus showed nuclear GFP localization as early as 24 h after infection and the proportion of nuclear signal increased at later time points (Figure 3A). Reporter cells infected with DENV-faR showed a similar trend, although a lower number of reporting cells was observed at 24 h and 48 h post infection in comparison to WT infected reporter cells (Figure 3A), consistent with lower replication capacity of the reporter virus (28). Cells infected with DENV-faR showed an increase of red fluorescence in a time-dependent manner providing evidence for viral replication and spread. Importantly, ∼83% to 100% of cells exhibiting red fluorescence also showed nuclear translocation of GFP at 48 h or 72 h post infection, respectively (Figure 3A). Additionally, only 2-3% of cells were positive for nuclear GFP in absence of reporter virus signal. These results indicate that the reporter cell line successfully detected DENV infected cells and that the infected cells can be reliably identified as early as 24 h post infection. Furthermore, this experiment demonstrated the suitability of the reporter construct to define the percentage of DENV infected cells without the need for intracellular staining.

### Live cell imaging of cells expressing the DENV polyprotein

Recently, a plasmid-based expression system for induction of DENV replication organelles in transfected cells has been described (40). This system, designated “plasmid-induced replication organelle - dengue (pIRO-D), encodes the viral polyprotein that is translated from an RNA generated in the cytoplasm by a stably expressed T7 RNA polymerase. In this way, the pIRO-D system allows the analysis of viral proteins in cells, independent of viral replication. However, since no fluorescent protein coding sequence is incorporated into the construct, expression of the DENV polyprotein cannot be followed by live cell imaging.

To overcome this limitation, we determined whether our DENV reporter cell line could be combined with the pIRO-D system to analyze the expression of the viral polyprotein in real-time. Huh7-Lunet cells stably expressing the T7 RNA polymerase and the reporter construct (Lunet-T7-RC) were seeded in dishes with glass bottoms and on the next day transfected with the pIRO-D construct (Figure 3B). The growth medium was changed to imaging medium at 4 h after transfection and the dishes transferred to a live cell imaging microscope. The GFP signal was recorded every 10 min for 8 h (final time point at 12 h post transfection). Representative images of mock transfected and pIRO-D-transfected cells at 2 h increments are shown in Figure 3C; a video spanning an 8 h observation period of transfected cells is shown in supplementary movie S1.

No nuclear translocation of the GFP reporter was detected in mock transfected Lunet-T7-RC cells (Figure 3C, upper row). In pIRO-D-transfected cells, nuclear localization was already detected in a few cells as early as 4 h post transfection, suggesting a robust expression of the viral polyprotein. The number of cells with nuclear GFP signal as well as the intensity of the signal increased over time.

### Development of a reporter system for identification of SARS-CoV-2 infected cells

The ability to rapidly detect virus infection in cell culture on a large scale is a valuable tool for studies of virus-host interactions, but also for identification and evaluation of antiviral drugs. The recent outbreak of SARS-CoV-2 has created a dire need for such tools to characterize virus infection and develop therapeutics. Therefore, we adapted and optimized the reporter system to sense SARS-CoV-2 infection. The first two open reading frames of coronaviruses are expressed as polyproteins ORF1a/b, which are cleaved into the individual proteins by viral proteases PL_pro_ and 3CL_pro_ (14–16). The sequence of the SARS-CoV-1 Frankfurt isolate was analyzed to determine the protease cleavage sites between individual nsps, and the deduced sequences were inserted into the linker region of our reporter construct (Table 4). The generation of constructs 2, 11 and 13 failed and they were not further pursued.

**TABLE 4.**
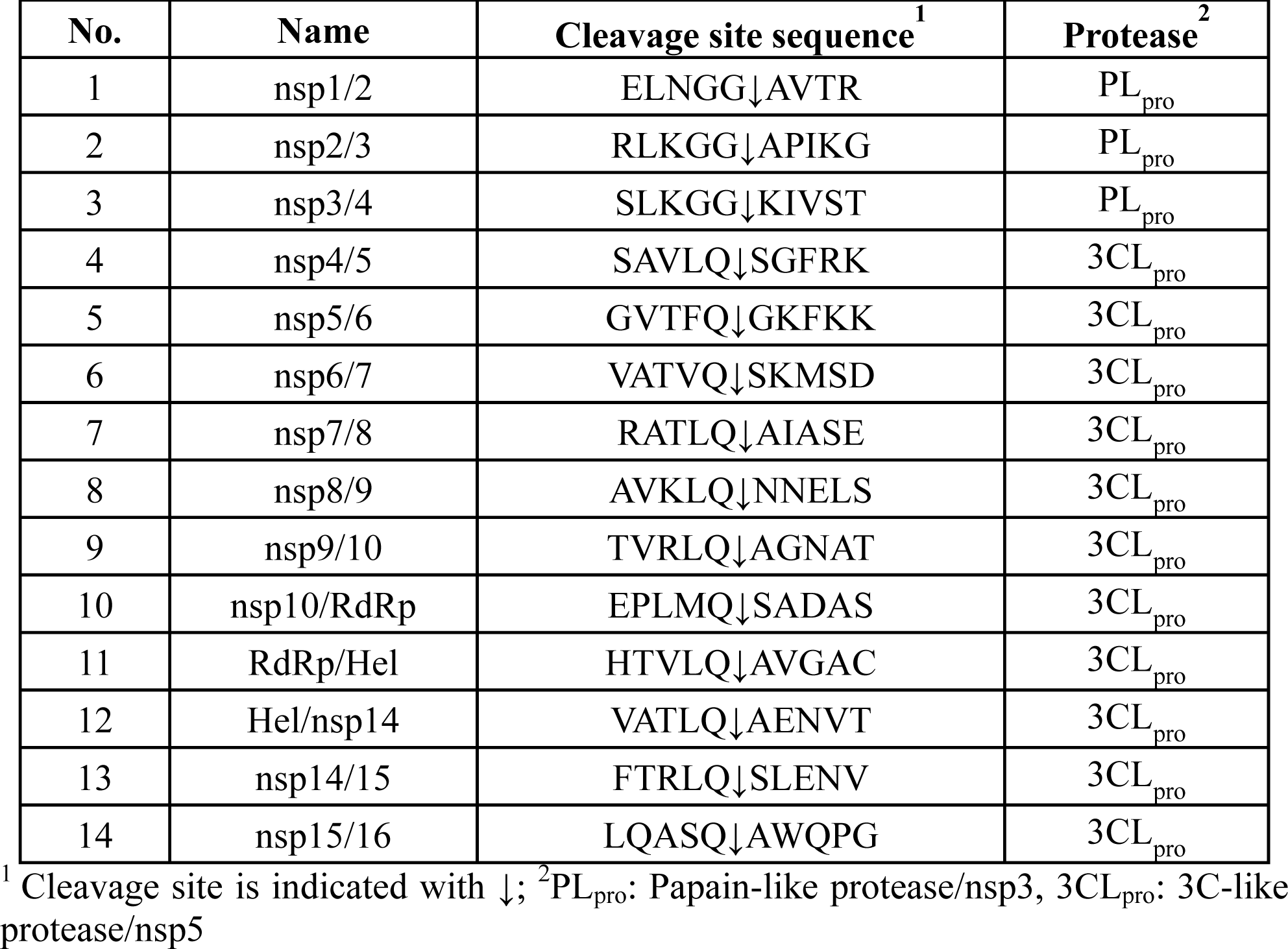
List of cleavage site sequences inserted into the SARS-CoV-2 reporter construct.

Cells expressing the GFP reporter containing the individual cleavage site linkers were generated by lentiviral transduction of A549 cells stably expressing the SARS-CoV-2 receptor ACE2 (A549-ACE2) (41). Productive infection of the cells with SARS-CoV-2 (strain BavPat1) was determined by detection of double-stranded (ds) RNA 48 h post infection. Cytosolic GFP localization was observed in all mock infected cells expressing the reporter constructs except for construct 3, where nuclear signal was evident (Figure 4A and 4B). No clear differences in GFP localization between mock and virus infected cells were observed for reporter constructs 1, 5 and 7 (Figure 4C), while all the others showed an increase in nuclear GFP signal upon infection (Figure 4C and 4D). Among the different constructs, the reporter construct 14 was the most sensitive, showing the highest number of cells double positive for dsRNA and nuclear GFP together with the lowest number of cells single positive for dsRNA staining (Figure 4D) and therefore was selected for further investigation.

**Figure 4:**
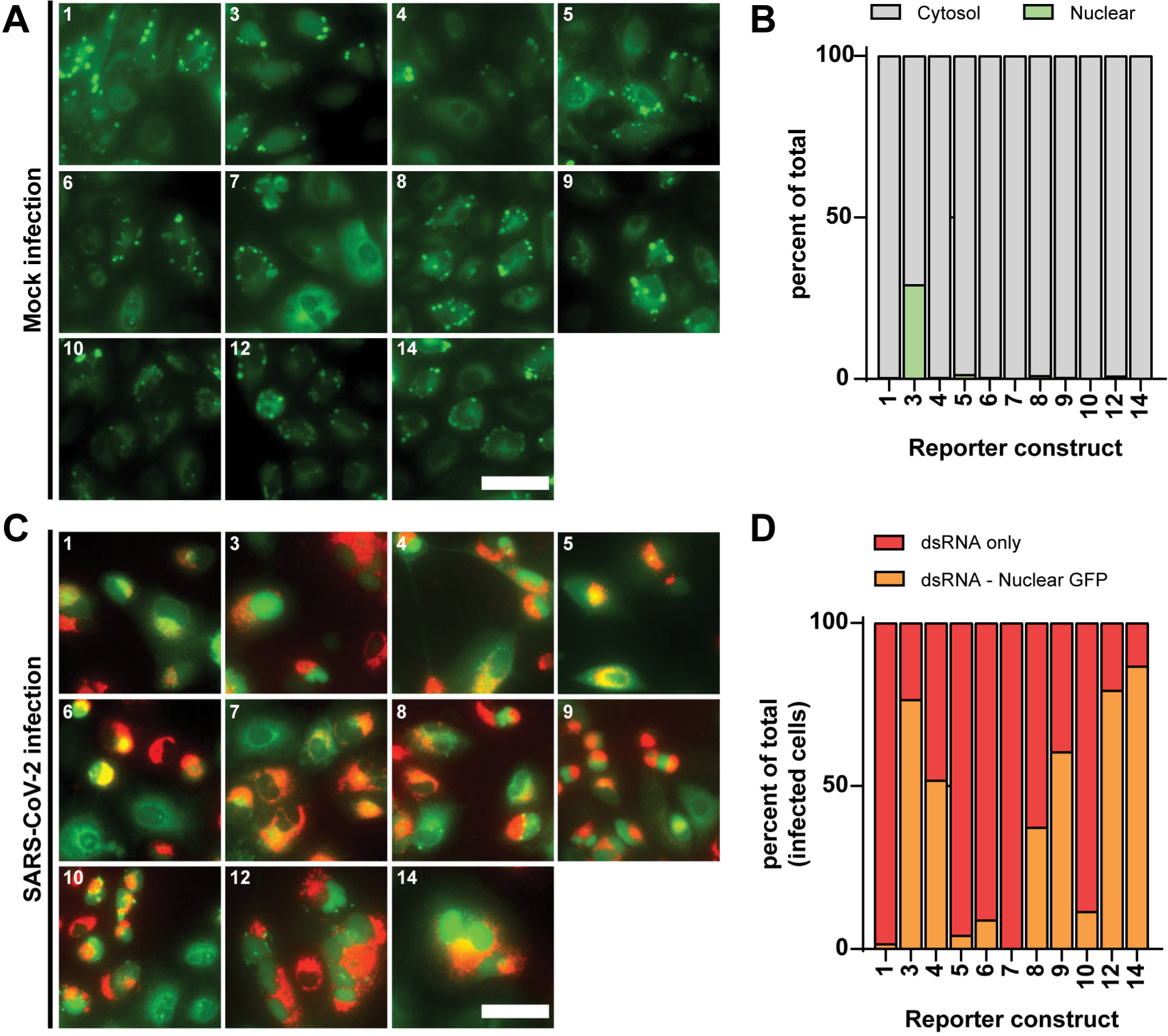
Screening of SARS-CoV-2 reporter constructs. A) A549-ACE2 cells were transduced with lentiviruses encoding for the reporter constructs specified on the top left of each panel. Cells were fixed 32 hours post-transduction and GFP localization was analyzed using a wide-field fluorescence microscope. B) Quantification of images acquired as in A). The percentage of nuclear or cytosolic GFP is shown (light green and gray, respectively). C) Cells transduced as in A) were infected after 16 hour post-transduction with SARS-CoV-2 (MOI = 5). Cells were fixed 16 hpi and viral double-stranded RNA (red), a replication intermediate indicative of active viral replication, and the GFP-based reporter construct (green) were detected by immunofluorescence using a wide-field fluorescence microscope. D) Quantification of images acquired as in c). Percentage of infected cells positive for dsRNA only (red) or double positive for nuclear GFP signal and dsRNA (orange) is shown. Scale bars: 50 μm.

### Live cell imaging of SARS-CoV-2 infection

To determine the kinetics of the reporting activity we investigated SARS-CoV-2 infection in reporter cells by live cell imaging. Since transiently transduced cells expressing the SARS-CoV-2 reporter construct showed highly heterogenous GFP signal intensity as well as the formation of large fluorescent aggregates (Figures 4A and 4C), we generated single cell clones by FACS sorting for cells with low expression. Among the twenty cell clones generated, clone C2 was selected based both on SARS-CoV-2 infection susceptibility and reporting activity (data not shown). The A549-ACE2 cells were transiently transduced with lentivirus encoding for the SARS-CoV-2 reporter construct 14 and seeded on glass bottom plates. After 24 h, cells were infected with SARS-CoV-2 (Figure 5A). Previous studies have determined that a complete virus replication cycle can occur within 6 h after infection but that virus replication and assembly continues to increase up to 24 h in A549-ACE2 cells (41). Therefore, live cell imaging was started at 2 h post infection and images were acquired every 10 min for 18 h (final time point 20 h post infection) (supplemental Movie S2). Representative images of mock and SARS-CoV-2 infected cells are shown in Figure 5B.

**Figure 5:**
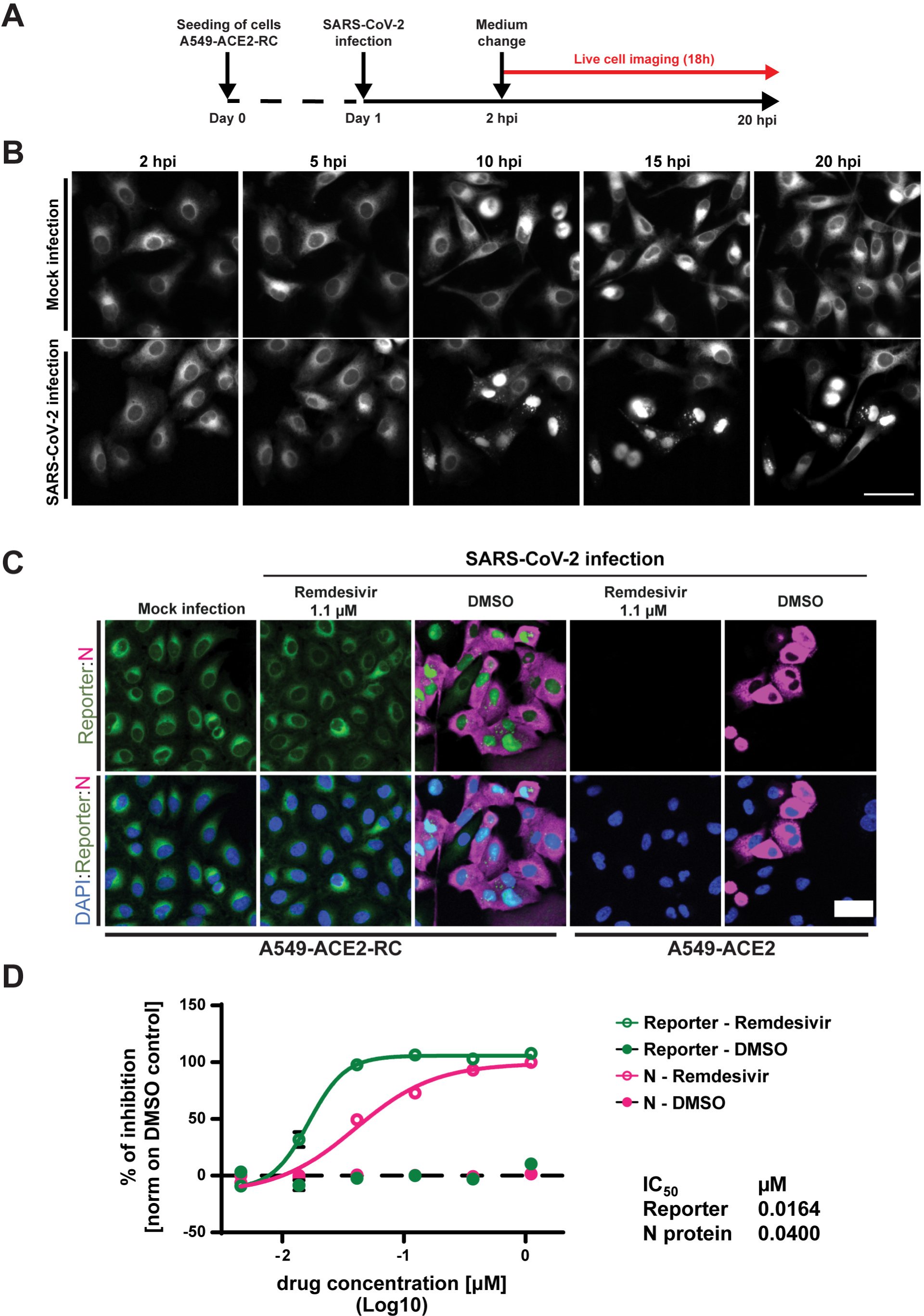
Application of the SARS-CoV-2 reporter cell line for live cell imaging of viral infection and assessment of antiviral activity of Remdesivir. A) Experimental set-up to monitor GFP-reporter activation in SARS-CoV-2 infected cells. B) A549-ACE2-RC (clone C2) cells stably expressing the reporter construct 14 were infected with SARS-CoV-2 (MOI = 10). 2 hpi live cell imaging was performed for 18 h using a confocal spinning disc microscope. Images of representative fields of view and time points are displayed. Scale bar: 50 μm. C) A549-ACE2 and A549-ACE2-RC (clone C2) expressing the SARS-CoV-2 reporter construct 14 were incubated with Remdesivir (1.1 μM) or DMSO control and infected with SARS-CoV-2 (MOI = 5). After 16 h, cells were fixed and stained for N protein before imaging with a confocal spinning disc microscope. Scale bar: 50 μm. D) IC50 calculation of Remdesivir in reporter cell clone 2 infected with SARS-CoV-2 (MOI = 5). Percentage of inhibition was calculated by quantification of the number of N-positive cells and cells with nuclear GFP signal in duplicate wells for each compound concentration. Values were normalized by setting the average number of infected cells in the DMSO treated sample as 0 % inhibition.

Mock infected cells exhibited an ER-like GFP signal throughout the observed time frame. In infected cells, a time-dependent increase in the numbers of cells showing nuclear GFP signal was observed with the earliest translocation event starting at 5.5 h post infection (supplementary Movie S2). These data demonstrate the suitability of our reporter system for live cell imaging analysis of SARS-CoV-2 infection.

### Application of the SARS-CoV-2 reporter for drug screening

The need for effective treatment for COVID-19 prompted us to investigate the suitability of the reporter cell line for drug screening. A proof-of-concept experiment was performed by using the nucleoside analogue Remdesivir, which is currently the only FDA-approved drug for treatment of SARS-CoV-2 infection. Both the A549-ACE2 C2 reporter clone and the parental A549-ACE2 cells without reporter construct expression were incubated with serial dilutions of Remdesivir for 30 min and infected with SARS-CoV-2 (MOI = 5). The compound remained present throughout the duration of the experiment. Cells were fixed at 16 h post infection and GFP translocation was evaluated using confocal microscopy.

Treatment with 1.1 μM Remdesivir lead to the loss of viral N protein fluorescence signal in both cell lines confirming the previously described antiviral activity (42, 43) (Figure 5C). Nuclear GFP signal was observed in DMSO treated reporter cells infected with SARS-CoV-2 confirming the reporting activity of the cell line. The percentage of cells displaying nuclear GFP signal and cytosolic N protein staining in the different Remdesivir dilutions was quantified by a semi-automated image analysis workflow (31) (Figure 5D). The IC_50_ calculated for the N protein staining was 40 nM (R^2^ = 0.987, 95% CI = 27.05 – 55.89 nM), in line with IC_50_ values found in our in-house assay (H. Kim and R. Bartenschlager, unpublished). Quantification with the reporter signal showed an IC_50_ of 14 nM (R^2^ = 0.997, 95% CI = 15.21 – 17.97 nM). This decrease in IC_50_ value is likely due to reduced sensitivity of the reporter construct that relies on 3CL_pro_ activity and the higher background in the read-out in comparison to IF staining of the highly abundant N protein. Nevertheless, the two IC_50_ values differ by only a factor of ∼3 and therefore are quite comparable. Thus, our reporter system can be reliably used in primary screens to screen e.g. large compound libraries.

## DISCUSSION

This study describes the generation and characterization of a fluorescence-based reporter system for detection of DENV and SARS-CoV-2 infection. The reporter construct contains three functional elements: a fluorescent protein fused to an NLS, the TM domain of sec61β for ER membrane anchoring and an exchangeable protease cleavage site cassette located within the linker region that connects the fluorescent protein and the ER anchor. This design allows easy adaptation of the reporter system to other viruses that encode for specific viral proteases, especially for other positive-stranded RNA viruses replicating in the cytoplasm. The high selectivity and specificity of the selected constructs, as shown by IF, western blot and live cell imaging, render this tool suitable for applications that require single cell analysis, such as live cell imaging and correlative light-electron microscopy (CLEM) approaches. Moreover, combining our reporter cell lines with image analysis pipelines that quantify nuclear translocation events allows for rapid and robust assessment of antiviral efficacy of compounds or other antiviral interventions, including high-throughput screening of large compound libraries.

Among the different cleavage sites tested for DENV, the reporter construct 1, composed of the cleavage site between capsid and prM, allowed reliable and selective identification of infected or transfected cells with a construct expressing the viral polyprotein (Figures 2 and 3). Interestingly, for several DENV reporter constructs, nuclear GFP localization was observed in absence of the viral protease (Figure 2A). A possible explanation for this could be that unspecific cleavage of the linker region might be mediated by cellular proteases due to high levels of expression of the reporter construct upon transient transduction.

Reporter constructs for detection of flavivirus infection have been described previously were either cytosolic or employed viral non-structural proteins as ER anchors (21–23, 38). Most of these reporter systems rely on the expression of large fragments of viral proteins (21–23) which can alter the physiological stoichiometry of the viral proteins and induce undesired pleiotropic effects. Indeed, even expression of single NS proteins can affect cellular functions, such as alteration of mitochondrial morphodynamics by NS4B (44). In contrast, since our reporter construct does not contain viral sequences, except for the cleavage site, it is less prone to affect cellular pathways and processes.

The flexibility granted by the modular nature of our constructs allows for simple adaptation of the reporter system to different viruses that encode proteases acting in close proximity of the ER membranes. This allowed us to rapidly adapt the system to the detection of SARS-CoV-2 infection in cell culture. Transient transduction of cells with lentiviruses coding for the different constructs allowed fast screening and identification of the most suitable cleavage site (Figure 4). Notably, while it was sufficient to use cell pools under antibiotic selection for DENV reporter cell lines Huh7-RC and Lunet-T7-RC, we had to establish single cell clones for the SARS-CoV-2 reporter proteins because in most cells large fluorescent aggregates were observed (Figure 4A). This is likely due to differences in the ability of cell lines to respond to high levels of GFP fusion proteins. In addition to sorting for cells with lower reporter expression as done here, this problem might be overcome by employing less active promoter or by using an alternative fluorescent protein.

Live cell imaging demonstrated that SARS-CoV-2 infected cells can be identified as early as 5.5 h post infection (Figure 5B). Real-time identification of SARS-CoV-2-infected cells is currently mainly performed by employing recombinant viruses expressing reporter genes (45–47). While these studies report robust and reliable identification of infected cells, our reporter cell line has advantages in certain settings. The use of reporter viruses requires molecular clones and the adaptation of the genomic sequence for each different isolate, which for viruses with large RNA genomes like SARS-CoV-2 involves substantial cloning efforts. Additionally, the relatively high mutation and recombination rate of RNA viruses during genome replication makes reporter viruses inherently unstable. Moreover, integration of a reporter gene into the recombinant virus may attenuate the replication efficiency, as observed when comparing WT to DENV-faR virus infection (Figure 3B). In contrast, our cell line allows the detection of wildtype virus isolates, although in this system conserved cleavage site sequences are required.

We tested the suitability of the SARS-CoV-2 optimized reporter cell line to assess the antiviral activity of Remdesivir and determined an IC_50_ of 16 nM in our reporter system. By using N staining as an alternative read-out, we obtained a somewhat lower efficacy of Remdesivir (IC_50_ = 40 nM), which is closer to the data reported in the literature (42, 43, 48). The reduced sensitivity of the reporter construct, which relies on 3CL_pro_ activity and GFP translocation, might stem from failure to detect cells with low levels of infection whereas the highly expressed N protein is already detectable by IF. Additionally, the selected cell clone 2 might differ in Remdesivir metabolism which can impact the antiviral efficacy of the nucleoside analogue. It is important to note that the IC_50_ value varies depending on which cell lines and assays are employed and the IC_50_ values determined in this study are in a similar range with only a 3-fold difference. Nevertheless, compounds found to exhibit antiviral activity in our reporter construct should be further validated, e.g. by plaque assay and/or viral RNA level quantification. Therefore, the reporter cell line can be employed in a primary screen to lower the number of candidates for validation in more sophisticated and time-consuming assays, thus reducing costs and increasing speed.

Two additional applications of our system shall be mentioned. The first is the use for CLEM, a powerful imaging method that can be used to mark and identify cells of interest amongst a large number of non-infected or un-transfected cells and subsequent analysis of this traced cell by electron microscopy methods (49). Secondly, the reporter cell line as described here can also be employed for protease inhibitor screens in areas that do not have access to biosafety level 3 laboratories. This can be done by transient or stable expression of the protease individually and monitoring of the reduction of nuclear GFP as a result of protease inhibition, similar to a recent study (50).

In conclusion, we describe a reporter system suitable for the detection of DENV and SARS-CoV-2 infected cells. The system is easy to handle and flexible and should be applicable to any virus encoding a cytoplasmic protease. It is suitable for a large number of methods and assays, including high content screening. In all these respects, we expect this tool to facilitate investigations of virus-host interactions, but also the development of antiviral drugs that are urgently needed to curb pandemic viruses such as SARS-CoV-2.

## ACKNOWLEDGE

We are grateful to Monika Langlotz and to the ZMBH Flow Cytometry and FACS Core Facility (FFCF, Heidelberg, Germany) for sorting the A549-RC cells. We would like to acknowledge the microscopy support from the Infectious Diseases Imaging Platform (IDIP) at the Center for Integrative Infectious Disease Research, Heidelberg, Germany. This work was supported in part by the Deutsche Forschungsgemeinschaft (DFG, German Research Foundation) - Project number 240245660 – SFB 1129 (TP11 and TP13) to A.R. and R.B.

---

**Supplemental Movie S1**. Live cell imaging of Lunet-T7-RC cells transfected transfected with the pIRO-D system.

**Supplemental Movie S2**. Live cell imaging of A549-ACE2-RC (clone C2) cells infected with SARS-CoV-2

